# From Abandoned Scripts to FAIR Community Pipelines: Rescuing Orphan Bioinformatics Workflows with nf-core — Lessons from Light-Sheet Fluorescence Microscopy

**DOI:** 10.64898/2026.07.29.741447

**Authors:** Carolin Schwitalla, Luis Kuhn Cuellar, Matthias Hörtenhuber, Niklas Grote, Tatiana Woller, Irene Lamberti, Benjamin Pavie, Thomas Küstner, Felix A. Kyere, Ian Curtin, Jason L. Stein, Sven Nahnsen

## Abstract

**Background:** Research software is essential for modern data analysis but is often developed and maintained by a small number of researchers. When developers leave, software may become orphaned, limiting reuse and risking the loss of valuable domain knowledge and computational methods. While the FAIR Principles for Research Software (FAIR4RS) provide an essential foundation for improving the reuse of research software, compliance with these principles alone does not guarantee practical reusability. Here, we investigate whether orphaned scientific software can be systematically rescued and transformed into sustainable, reusable workflows using established software engineering practices and community standards.

**Findings:** We re-engineered the abandoned MATLAB-based NuMorph toolkit for large-scale light-sheet microscopy image analysis into nf-core/lsmquant, a Nextflow-based workflow developed according to nf-core community guidelines. The re-engineered workflow preserved the original scientific methods at comparable computational cost while improving the software’s FAIRness, portability, and reproducibility. Integration into the nf-core ecosystem provides a community-driven framework that supports software sustainability through distributed maintenance and shared development practices, while the modular workflow architecture simplified adaptation of nf-core/lsmquant to additional light-sheet microscopy datasets beyond the original application

**Conclusion:** Our work demonstrates that orphaned scientific software can be successfully rescued through systematic re-engineering guided by FAIR and software sustainability principles. By transforming a legacy codebase into a community-maintained workflow, we preserve valuable domain-specific methods while improving usability, maintainability, and reproducibility. This approach provides a practical strategy for recovering orphan research software and integrating it into modern, reusable research ecosystems.

## Introduction

Scientific research in the life sciences increasingly relies on specialized software and complex computational tasks to process large-scale omics and imaging datasets to extract biological insight. Computational workflows integrate independent tools to automate processing and analysis tasks and have emerged as an essential response to the rapid growth in data generation driven by modern experimental technologies. Despite the central role of research software (software developed during or for research purposes[1]) in enabling the analysis of rapidly growing datasets, the reuse of existing tools and reproducibility of workflows remain difficult. Commonly observed challenges for reusing research software and reproducing results are poor documentation, difficulties in installation and deployment, limited interoperability among compute infrastructures, the availability of code, insufficient description of parameter settings, and incomplete, outdated, or incompatible dependencies[2–6]. The underlying reasons for these issues are multifaceted. Research software is developed under academic conditions that prioritize innovation and publication over software quality and long-term maintainability. In contrast to industry, software development is often carried out by individuals or small teams, who rarely receive any formal training in software engineering practices[7,8]. When lead developers leave, software projects are often abandoned, remaining theoretically functional but practically unusable due to poor maintenance. This can result in the loss of domain-specific software tools and expertise, forcing researchers to reimplement methods rather than build on existing solutions. Consequently, this situation slows scientific progress and compromises reproducibility. The challenges associated with reproducibility and reuse of research output have received more attention over the years, and as a response, the FAIR (Findable, Accessible, Interoperable, and Reusable) principles for data stewardship and FAIR for research software (FAIR4RS) were introduced[1,9,10].

These guidelines aim to ensure that research output and software can be reliably discovered, cited, reused, and extended, thereby improving reproducibility and long-term usability[1,10,11].

Alongside the life sciences, the field of software engineering research has also responded to the challenges of software reuse, integration, and adaptation to technological progress by introducing the concept of *software sustainability*. The Karlskrona Manifesto for Sustainability Design describes software sustainability as a multi-dimensional property, including social, economic, environmental, as well as a technical dimension[12]. The last refers to software that remains functional over time, can be maintained, and can evolve with advancing technology and changing conditions[13]. More specifically, sustainability was proposed as a non-functional requirement for software, measured by the following properties: Extensibility, Interoperability, Maintainability, Portability, Reusability, Scalability, and Usability[14]. The concept is further promoted by the 2010-founded software sustainability institute, which advocates for the importance of research software, good software practices, and shaping the current understanding and reward system to acknowledge software as a valid research output[15]. Both frameworks, software sustainability and FAIR principles, go hand in hand, emphasizing the central role of reuse for knowledge growth, reproducibility, and transparency in scientific practice. Based on these conceptual frameworks, multiple recommendations have been proposed to translate these principles into practical development practices. For example, the use of version control systems to track code changes and software versions, as well as package managers like Conda[16] and containerization technologies, such as Docker[17] or Apptainer (formerly Singularity)[18], to manage software dependencies and computing environments[19,20]. In addition, workflow management systems like Common Workflow Language (CWL)[21], Nextflow[22], Snakemake[23], or Galaxy[24] are useful solutions to build end-to-end workflows running multiple different processing steps and tools in a controlled, reproducible, and scalable way[25]. Clear licensing, comprehensive documentation, and simple installation procedures facilitate software reuse. Community standards for data formats, metadata, and interfaces improve interoperability and workflow integration, while public repositories enable transparent sharing and reuse within the scientific community[26–28].

While these practices can be implemented by individual developers or research groups, coordinated community efforts have emerged to promote their broader adoption and to provide shared frameworks and standards for sustainable, FAIR research software development, maintenance, and sharing. One example is the *nf-core community*, founded in 2018 around the Nextflow workflow management system[29]. Pipelines developed within the nf-core ecosystem follow established software engineering practices, including version control, use of containerized software, code review, continuous integration, automated testing, and extensive documentation. Standardized installation procedures and simplified execution commands further facilitate adoption by users across diverse computational environments. Furthermore, nf-core promotes collaborative development through training activities, hackathons, and open community contributions, enabling distributed maintenance and knowledge exchange among researchers. With the combination of community standards and collaborative development, nf-core implements key principles of reproducibility and sustainable software development. A similar community-driven approach is the Galaxy platform, which provides an open framework for constructing and sharing reproducible computational workflows through an accessible interface[24].

These community frameworks have historically been concentrated in classical omics domains like genomics and transcriptomics, driven by the advancing next-generation sequencing technologies. Recently, bioimaging technology has become a rapidly emerging area of research through advances in high-throughput imaging, particularly light-sheet fluorescence microscopy. This technology enables the acquisition of whole-organ and organism images at cellular resolution, frequently generating data in the terabyte range. Processing such large datasets remains difficult. Existing tools, often based on a graphical user interface, frequently require substantial manual intervention, like custom scripting for automation, and barely scale. As a result, data processing and analysis increasingly represent a critical bottleneck for researchers. One notable effort to address the challenges of large-scale image processing is NuMorph[30], a toolkit for end-to-end processing of terabyte-scale whole-mouse brain light-sheet microscopy data. Despite its methodological relevance, NuMorph suffered from limited user adoption, and its maintenance and further development were discontinued after the developer left the research group where NuMorph was developed. Ultimately, the NuMorph toolkit became *orphan software*, a term that is used to describe code that has no maintainer and no activity for a long time[31,32]. In this work, we demonstrate a systematic approach for *rescuing orphan bioinformatics code* by re-engineering the legacy NuMorph toolkit into a sustainable workflow, nf-core/lsmquant[33]. We show how FAIR and software sustainability recommendations can be applied to convert such a legacy codebase into a reusable, community-maintained pipeline. Specifically, we use Nextflow, which enables substantial improvements in scalability, portability, and reproducibility due to its inherent process parallelization, compatibility with most widely used operating systems, workload managers, and cloud infrastructures, and the usage of containerized execution environments. Furthermore, community guidelines for documentation, workflow design, standardized interfaces, and continuous integration practices facilitate the practical implementation of FAIR and sustainable software properties. We show that the re-engineered workflow nf-core/lsmquant produces results identical to those generated by the original NuMorph toolkit, preserving previously established methods and benefits from Nextflow’s parallelization. Lastly, we apply nf-core/lsmquant to datasets other than the original use case of whole mouse brain images, demonstrating the potential to generalize to other applications. Taken together, our work shows a generalizable strategy for rescuing orphaned scientific software and integrating it into modern, reproducible research ecosystems, thereby preserving valuable domain knowledge and enabling broader adoption within the research community.

### Findings

The migration of the legacy system NuMorph to a Nextflow-based nf-core pipeline followed the general model for software re-engineering. The general model consists of three consecutive stages: reverse engineering, alteration, and forward engineering. During reverse engineering, information about the function, structure, dependencies, and design of the legacy system is recovered. In the alteration process, the legacy system components and source code are modified to match the target system’s specifications. Finally, within forward engineering, the target system is implemented[34,35].

We applied this procedure to the NuMorph toolkit, resulting in the pipeline nf-core/lsmquant. The migration wraps a tightly coupled, script-based implementation in a modular, containerized workflow that enables reproducible, scalable execution across computing environments. In the following, we first describe the re-engineering process of the migration. We then evaluate the resulting workflow with respect to scalability, reproducibility, and validity of results compared to the original implementation. Finally, we assess the extent to which the re-engineered pipeline adheres to FAIR and software sustainability principles and discuss the implications for reuse and long-term maintainability.

## 1. Re-engineering of NuMorph

### Reverse engineering

To assess NuMorph’s structure and functionality, we examined the toolkit’s source code, documentation, and publication.

NuMorph’s source code is publicly available on Bitbucket, GitHub, and Zenodo and is primarily implemented in MATLAB (231 files, 18,573 lines of code) with a smaller Python component (28 files, 2,314 lines of code). The repository on Bitbucket contains an incomplete wiki page for the software, describing requirements, basic setup, run commands, and sample information. Further sections of the wiki page are present but empty. The repository on GitHub contains a short readme describing the toolkit’s main functions and provides a link to the test dataset within a few sentences.

NuMorph’s implementation follows a procedural programming style with control flow realized via nested if–else blocks. It relies on a complex parameter configuration structure, which is adjustable by the user, using dedicated MATLAB scripts and is partially computed at runtime. For internal computations and output organization, NuMorph creates and depends on a specific folder structure. Altogether, the source code contains a complex coupling of function scripts, control flow, and environment, which complicates maintenance and reuse. Using the information from the source code, documentation, and publication, we derived an abstract representation of NuMorph’s process units and workflow (Fig.2).

**Figure 1:**
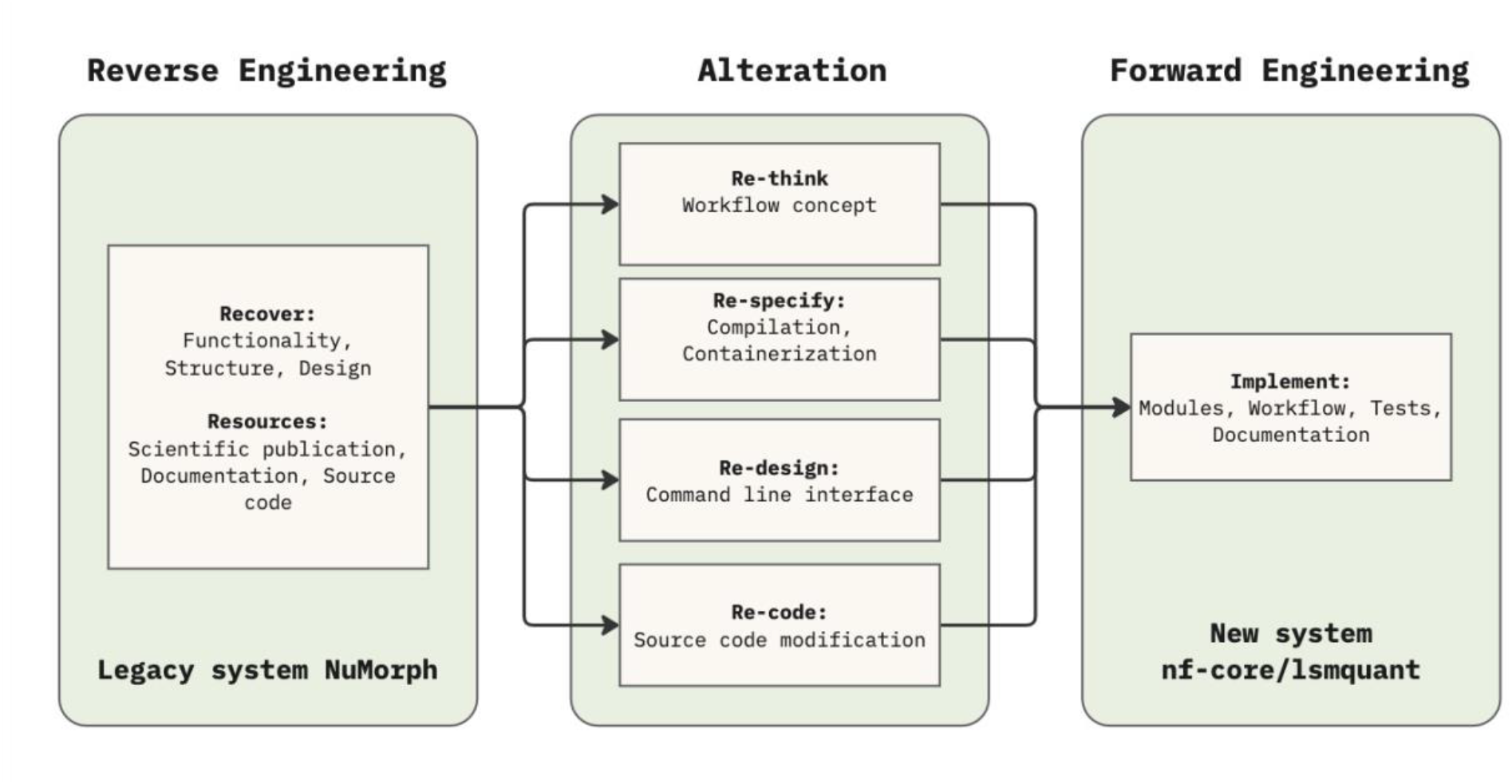
Re-engineering strategy for the migration of the legacy system NuMorph to the new system nf-core/lsmquant. Schematic overview of the three-phase re-engineering approach, adapted from the general model for re-engineering, used to migrate the legacy NuMorph system to the nf-core/lsmquant pipeline. In the Reverse Engineering phase, the existing NuMorph system was analyzed by recovering its functionality, structure, and design from available resources, including scientific publications, documentation, and source code. The recovered knowledge informed four parallel Alteration steps: (1) *Re-think*, reconceptualizing the overall workflow design; (2) *Re-specify*, addressing compilation and containerization; (3) *Re-design*, adapting the command line interface; and (4) *Re-code*, modifying the underlying source code. The outcomes of these alteration steps were integrated in the Forward Engineering phase, where the new nf-core/lsmquant pipeline was implemented, encompassing Nextflow modules, the workflow structure, test infrastructure, and documentation.

**Figure 2:**
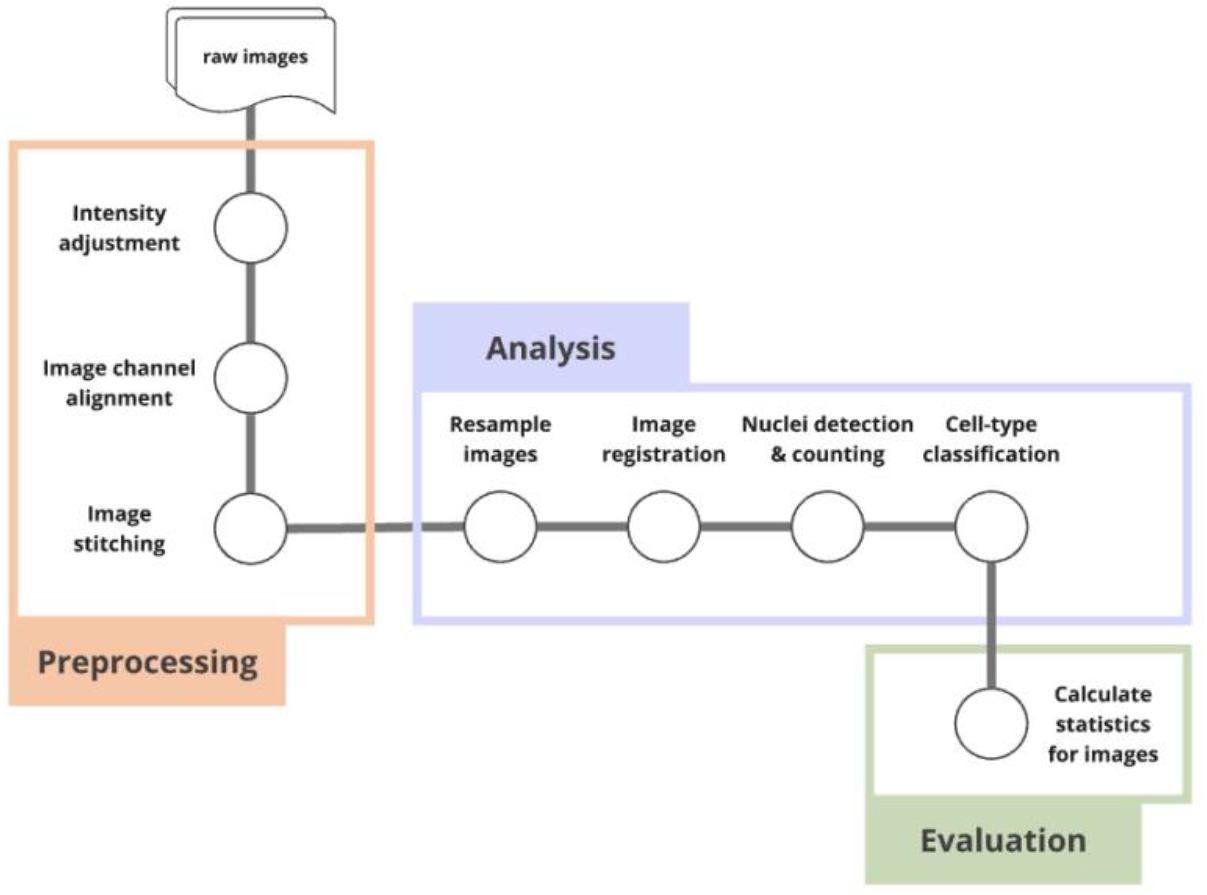
Abstract representation of the NuMorph workflow. Logical processing units are depicted as circles connected by a grey line representing the common dataflow through the system. The workflow is organized into three stages: Preprocessing, which takes raw images as input and sequentially performs intensity adjustment, image channel alignment, and image stitching; Analysis, which encompasses resampling, image registration, nuclei detection and counting, and cell-type classification; and Evaluation, which calculates summary statistics for the processed images.

Conceptually, the toolkit is organized into three main stages: preprocessing, analysis, and evaluation, each containing logically separable processes.

We focused on three core preprocessing steps (intensity adjustment, channel alignment, iterative stitching) and two key analysis processes (registration to the Allen reference brain atlas and cell nuclei quantification using a 3D U-Net[36], identifying these five processes as primary candidates for implementation as Nextflow modules.

### Alteration

Based on the information recovered from reverse engineering NuMorph, we identified multiple necessary modifications to the original system. We modified the source code and implemented a command-line interface (CLI) to enable parameter configuration via the open comma-separated values (CSV) file format, controlling the execution of NuMorph processes and combining individual start-up steps into a single command. For distribution, the NuMorph stages: *preprocessing* and *analysis,* were compiled and containerized via Docker to package the candidate processes: *intensity adjustment*, *channel alignment*, *iterative stitching*, and *atlas registration*. The Python-based 3D U-Net *cell nuclei quantification* was separated from the MATLAB codebase, restructured into a pip installable package, and a CLI was added to enable execution and parse process-specific parameters. Due to outdated and incompatible dependencies in the originally provided Conda environment, dependencies were updated and revised. A corresponding Docker image was then created to package the updated Python-based segmentation.

### Forward engineering of nf-core/lsmquant

By using the containerized functions of NuMorph from the alteration process, we implemented nf-core/lsmquant. The workflow was implemented in Nextflow domain-specific language 2 (DSL2), the nf-core pipeline template, and following community best practices. Each of the five target processes that we identified during reverse engineering was implemented as a Nextflow module and executes NuMorph’s containerized processes as independent units, orchestrated by Nextflow.

### Input specification

The pipeline input is specified using a samplesheet CSV file, containing three columns per sample. The first column contains a user-defined sample identifier. The second column provides the path to the raw image data, either as a ZIP archive or a directory. The pipeline automatically detects the input type and stages the data into the Nextflow working directory using either the *stagefiles* module for input directories or the nf-core *unzip* module for compressed archives. The raw light-sheet data are required to be 16-bit 2D TIFF images, organized as one image per channel, per tile, and per z-slice. The third column specifies the path to a parameter CSV file containing processing-specific configuration parameters. This file enables users to adjust individual processing steps and is passed to the NuMorph tools to reconstruct the global configuration during execution. A template for the parameter CSV file is provided within the pipeline repository. To ensure correct parameter declaration in the parameter CSV file, we implemented a JSON schema, which is used for parameter validation in the module *validate_parameters*.

### Workflow overview

Nf-core/lsmquant provides three workflows for processing raw TIFF images, which can be set via the parameter *stage*:

1. **stitch_only:** performs iterative tile stitching on raw images
2. **int_stitch:** Performs intensity adjustment and iterative tile stitching on raw images (default)
3. **int_align_stitch:** Performs intensity adjustment, channel alignment, and iterative stitching on raw images.

This design enables flexible processing of raw images, extending to a broader range of datasets, including single-channel datasets. For light-sheet microscopy data, iterative tile stitching is implemented as a mandatory processing step to merge individual tiles and reconstruct the complete image volume. To ensure generalizability, Allen Reference Atlas (ARA) registration and cell nuclei quantification were implemented as optional downstream analysis steps that can be set through the pipeline parameters *`--quantification`* and *`--ara-registration`*. Making these steps optional allows the pipeline to consider different use cases and needs while avoiding unnecessary computation. Each processing step generates intermediate results, which are stored in proprietary MATLAB (.mat) files in the original implementation. To provide user access to intermediate results without the need for a proprietary MATLAB license, we implemented a MATLAB-based conversion tool, *mat2json*, which converts these outputs to open CSV or JSON format.

### Documentation

In accordance with nf-core requirements, comprehensive documentation was added to describe pipeline usage and output. The nf-core template provides a standardized documentation structure, including general instructions for pipeline execution with Nextflow and consistent guidance across all nf-core pipelines. Additional sections were added to document the parameters specified in the CSV file, detailing their functions and impacts on individual processing steps. Furthermore, an extended method description was added to explain the functionality of each NuMorph process used in the pipeline. This information was obtained during the reverse engineering phase and extracted from the original NuMorph publication and source code.

## 2. Benchmark experiment

We conducted a benchmark experiment to evaluate resource usage and functional equivalence of nf-core/lsmquant and NuMorph. Each workflow was executed 30 times with the same parameter setting on the small test dataset from the original NuMorph toolkit, to record resource usage and workflow outputs for comparison. The small test dataset contains a subset of a tissue-cleared mouse brain, taken from z-plane 0600 to 0620 of 4 tiles and two channels: ctip2 (for lower layer excitatory neurons) and topro (TOPRO for all nuclei). In total, the small dataset consists of 169 images and has a size of 1.39 GiB

Both workflows executed their main workflow: intensity adjustment, channel alignment, stitching, and cell nuclei quantification. The experiments were performed on a Linux-based workstation with 16 CPU cores and 64 gigabytes (GB) of RAM, and one GPU (NVIDIA GeForce RTX 3090).

### Functional equivalence evaluation

To validate the functional equivalence of the new system nf-core/lsmquant to the legacy system NuMorph, we compared the outputs obtained by the benchmark experiment using the small test dataset. We calculated all pairwise mean square errors (MSE) for the final stitched volume of all 30 runs of the original NuMorph toolkit and nf-core/lsmquant for both channels and visualized the middle z slice for a random replicate of both workflows.

Figure 4B shows that pairwise MSE calculations across the 30 workflow runs of NuMorph and nf- core/lsmquant resulted in 0.0, meaning no voxel differences were observed between the two workflows. As a qualitative example, we visualized the middle z-plane (z 0009) of randomly picked replicates from each workflow and their corresponding intensity difference map (Figure 4A). This shows that the reengineering procedure of the legacy system NuMorph did not introduce changes that influence the final stitched images. Since the calculated intensity adjustments and alignment transformations are applied during the stitching process, the preprocessing stage of nf- core/lsmquant, including all three processes: intensity adjustment, channel alignment, and stitching, is functional equivalent to the legacy system NuMorph using the small test dataset.

**Figure 3:**
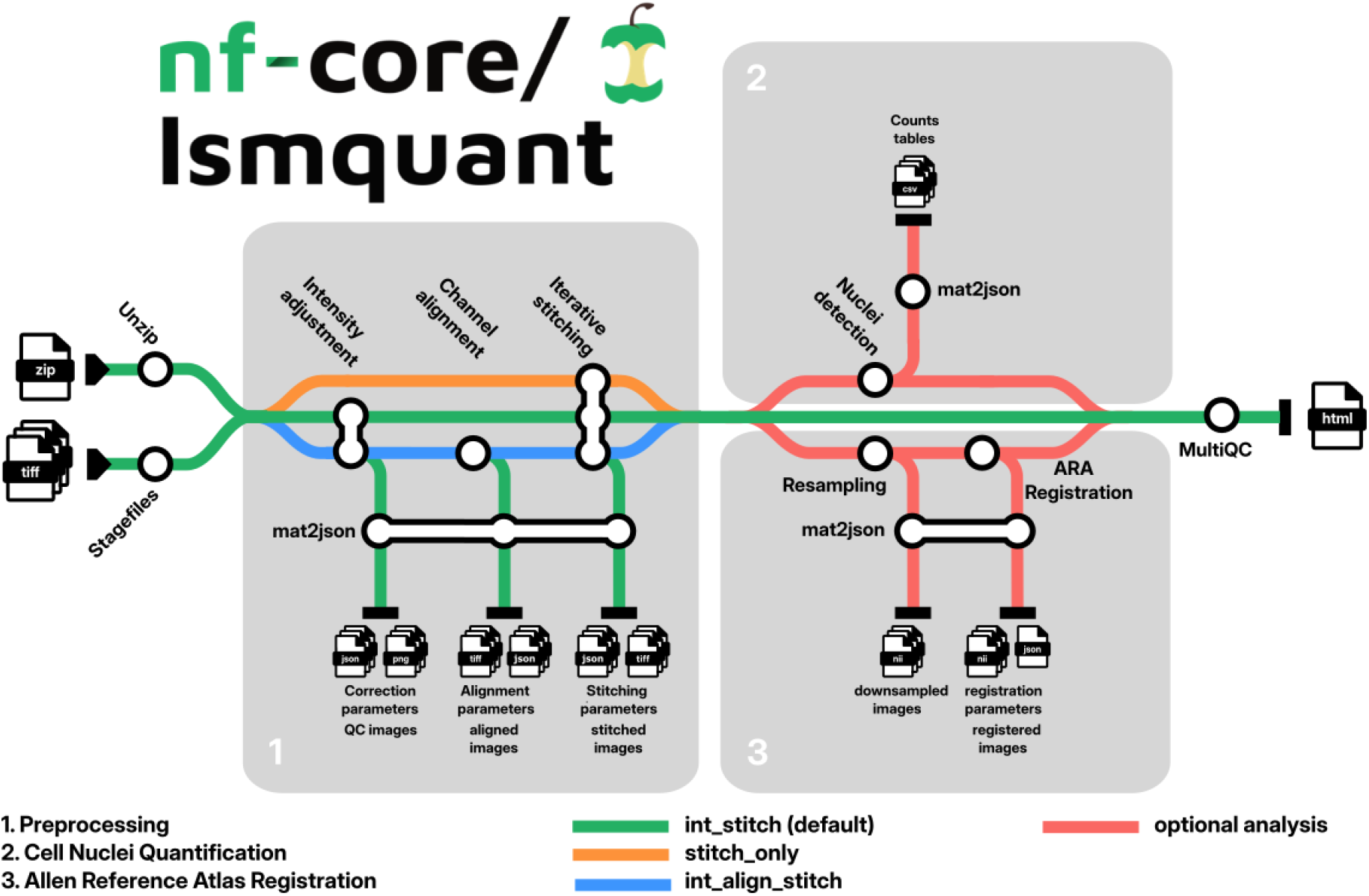
Abstract representation of nf-core/lsmquant as a metro map. The metro map illustrates the individual processes (circles) grouped into three logical processing stages. Input files (compressed archives and stage files) are first unzipped and passed into stage **1 (Preprocessing)**, which encompasses intensity adjustment, channel alignment, and iterative stitching. Intermediate outputs, including correction parameters, QC images, alignment parameters, and stitched images, are written to disk via mat2json. The pipeline offers three entry points: the default *int_stitch* mode (green), which executes the full preprocessing cascade; *stitch_only* (orange), which bypasses intensity adjustment and alignment; and *int_align_stitch* (blue), which performs intensity adjustment and alignment before stitching. Optionally, preprocessed images can be forwarded to **stage 2 (Cell Nuclei Quantification),** where nuclei detection and mat2json conversion produce count tables. Independently and optionally, **stage 3 (Allen Reference Atlas Registration)** performs resampling and atlas registration, generating down-sampled images, registration parameters, and registered images.

**Figure 4:**
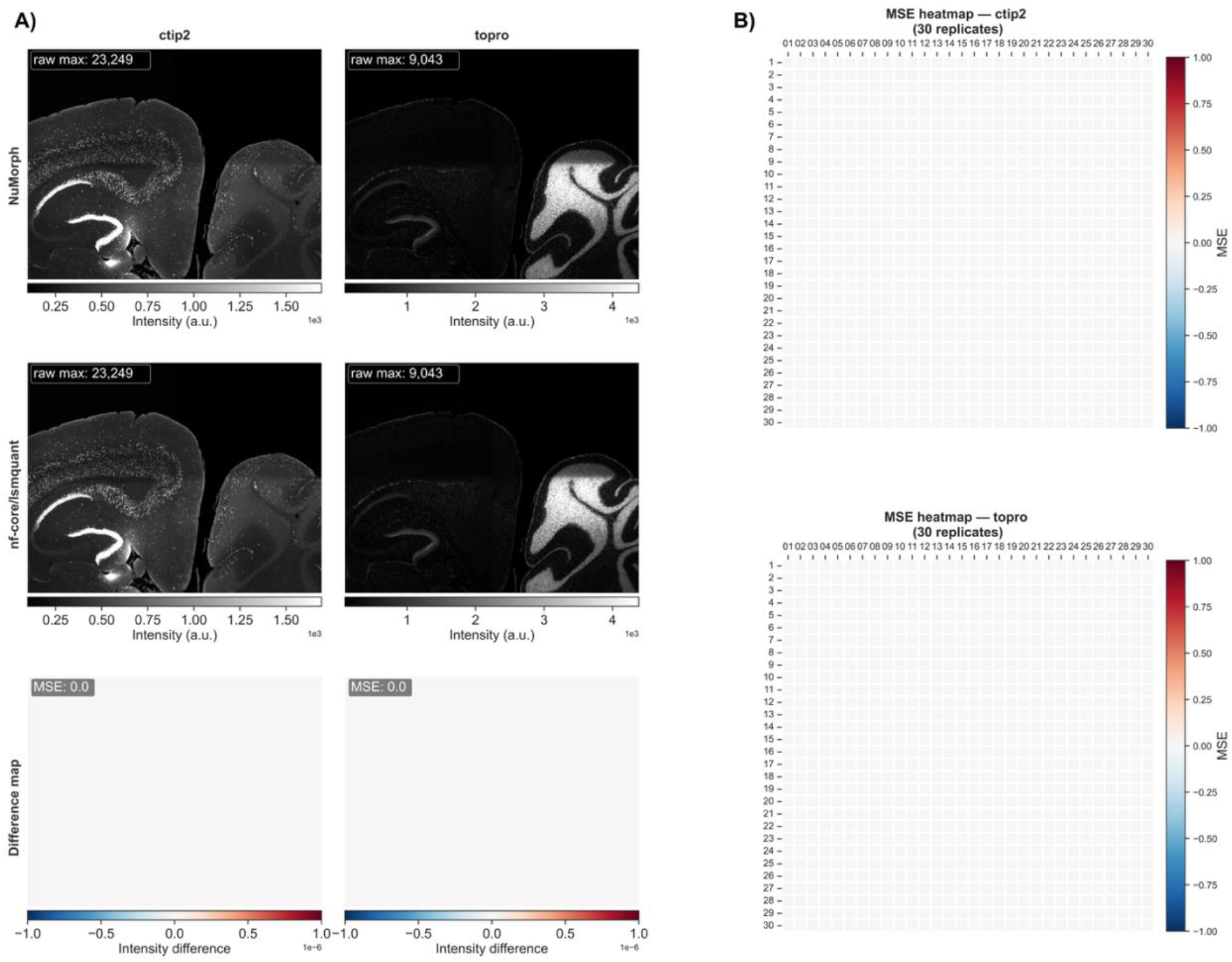
Comparison of stitched image outputs between the legacy system NuMorph and the new system nf-core/lsmquant. Representative stitched images from z-plane 0009 are shown for the channels ctip2 and topro. Grayscale images are contrast- stretched to the 1st–99th percentile of the NuMorph output, and the same intensity limits are applied to the corresponding nf-core/lsmquant image, to visualize relative brightness differences. For each channel, outputs generated by NuMorph and nf-core/lsmquant are compared using pixel-wise difference maps, with the corresponding mean squared error (MSE) reported for the displayed slice (A). Pairwise MSE heatmaps summarize comparisons of the complete stitched volumes across 30 independent pipeline runs performed using the same input dataset and identical parameter settings. Rows and columns correspond to individual runs. An MSE of 0.0 across all pairwise comparisons demonstrates that nf-core/lsmquant reproduces the stitching results of the original NuMorph implementation (B).

We also tested functional equivalence on a full-mouse brain dataset[37,38] containing 3 channels. We ran the preprocessing stage (intensity adjustment, channel alignment, and stitching) of each workflow on the same dataset and identical parameter settings.

Here, all MSE per z-plane of each channel resulted in 0.0 (Figure S1), and the visualized examples of the z-plane 0507 (Figure 5) show no pixel differences between the two workflows. This demonstrates that, also on a whole-mouse brain dataset, nf-core/lsmquant preserved NuMorph’s functionality.

**Figure 5:**
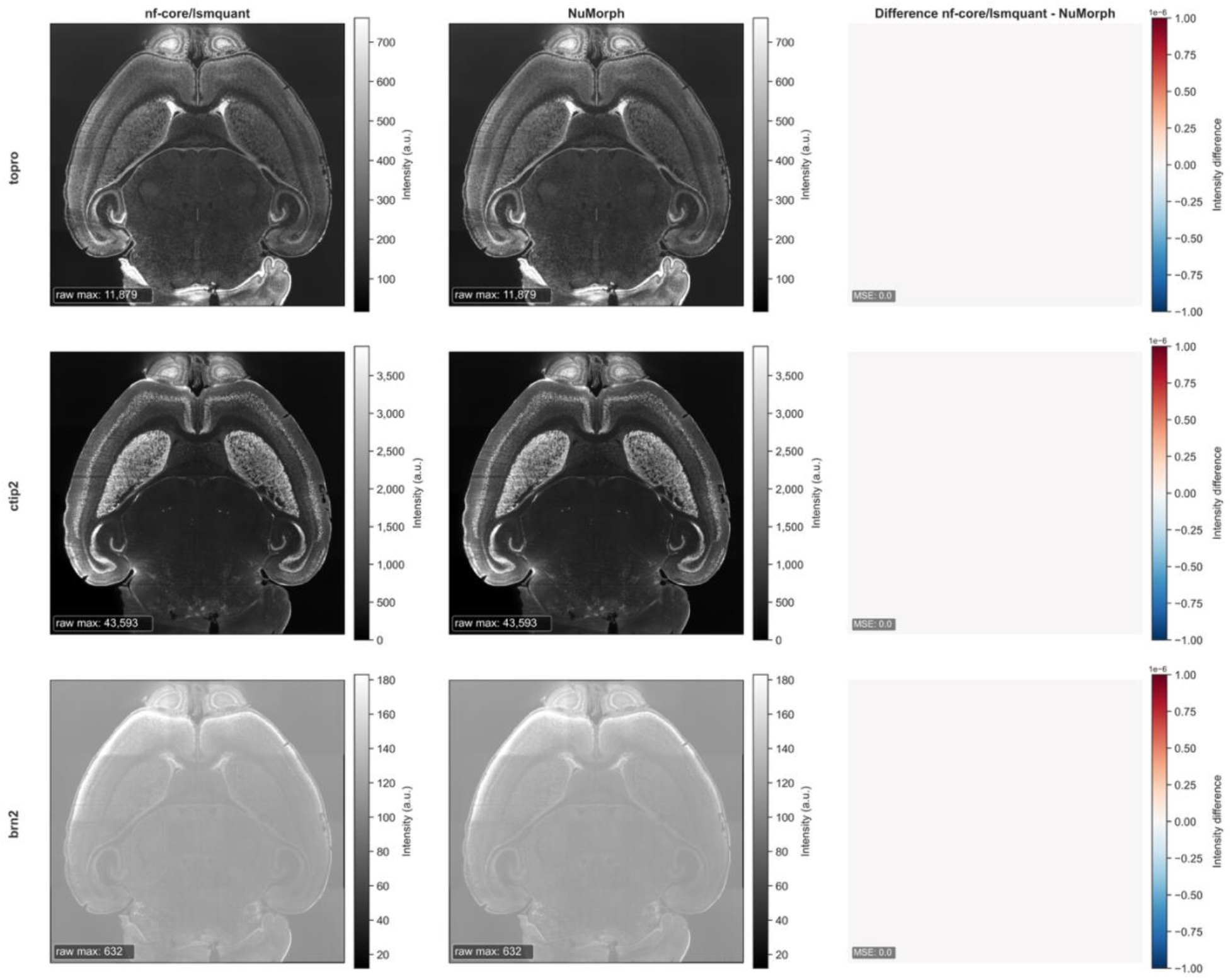
Comparison of stitched full mouse brain image outputs between the legacy system NuMorph and the new system nf-core/lsmquant. Representative stitched images from the middle z-plane 0507 are shown for the channels: topro, ctip2, brn2. Grayscale images are contrast stretched to the 1st–99th percentile of the NuMorph output, and the same intensity limits are applied to the corresponding nf-core/lsmquant image to visualize relative brightness differences. For each channel, outputs generated by NuMorph and nf-core/lsmquant are compared using pixel-wise difference maps, with the corresponding mean squared error (MSE) reported for the displayed slice.

**Figure 6:**
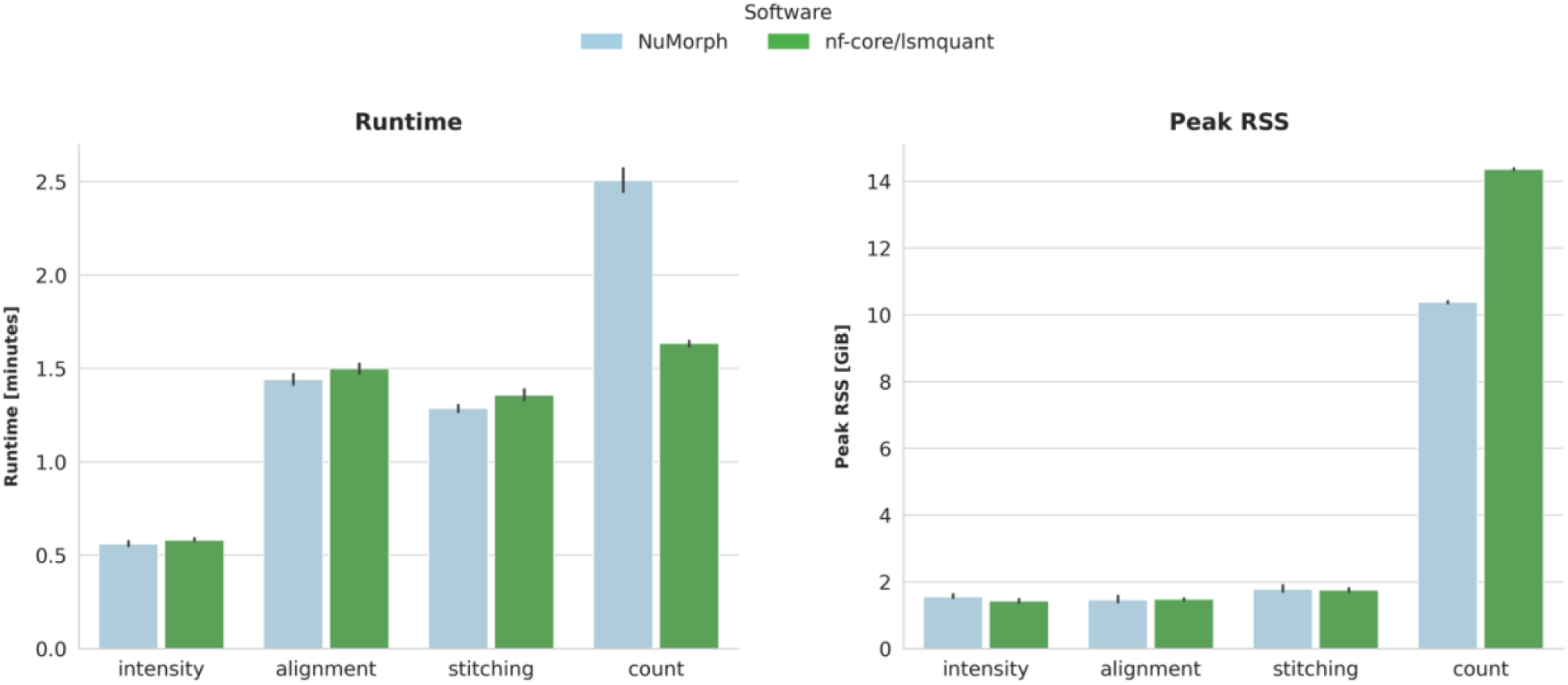
Computational performance comparison between NuMorph and nf-core/lsmquant across processing steps. Mean wall-clock runtime and peak resident set size (RSS) representing the maximum physical RAM are shown for each of the four processing steps: intensity adjustment, channel alignment, stitching, and cell nuclei counting. The bar plot heights indicate mean runtime and memory consumption per process, and error bars display the 95% interval of the data (ranging from 2.5 to the 97.5 percentiles).

### Performance evaluation

We compared the performance of nf-core/lsmquant with the legacy system NuMorph by tracking resource usage and runtimes for the processes: intensity adjustment, channel alignment, iterative stitching, and cell nuclei quantification during the benchmark experiment on the small test dataset

Runtime comparisons using the small test dataset show similar or slightly longer runtimes for nf-core/lsmquant for the processes: intensity adjustment, channel alignment, and stitching (Fig.5), where the mean runtimes differ between 1.2 and 4.2 seconds (Table S1). The marginally longer runtimes observed for nf-core/lsmquant could be caused by the start-up overhead inherent to per-process container execution. Also, the memory usage of these three processes is very similar between nf-core/lsmquant and NuMorph, with mean peak resident set size (RSS) differences between 18 and 123 megabytes (MiB). Here, only channel alignment of the nf-core/lsmquant implementation shows a slightly higher mean memory usage (∼18 MiB mean difference) than the legacy NuMorph implementation, where the other two processes consume slightly less compared to NuMorph (∼123 and ∼28 MiB mean difference).

In contrast, the cell nuclei quantification process (count), which is based on a trained 3D U-Net for centroid prediction, performs very differently in the two implementations (Figure 5). The count process in the nf-core/lsmquant implementation is substantially faster compared to the original NuMorph implementation, with a mean difference of ∼ 52 seconds (Table S1), but consumes ∼ 4 GiB (Table S1) more physical memory.

This difference in resource usage can most likely be explained by how we containerized the process. We used an NVIDIA-optimized base image to create the final Docker image for our reimplementation of the NuMorph 3D U-Net. This base image has a set of pre-installed ML libraries, Cuda and, a Nvidia optimized version of the original TensorFlow package for NVIDIA GPU architectures.

### Batch processing scalability

We evaluate the scalability of batch processing multiple samples between the new system nf-core/lsmquant and the legacy system NuMorph, by comparing the total batch processing runtime (wall-clock time) for 1 up to 10 samples.

The new implementation nf-core/lsmquant shows consistently lower runtime for batch processing 2 – 10 samples compared to NuMorph. Only when processing 1 sample NuMorph is slightly faster. This observation reflects the default execution model of the two systems. While native NuMorph processes multiple samples sequentially, nf-core/lsmquant leverages the task orchestration capabilities of the Nextflow workflow manager to execute independent tasks in parallel over multiple samples.

## 3. Real-world deployment and adoption

The development of nf-core/lsmquant within the nf-core ecosystem also facilitated collaboration beyond the original development group. Through community interactions, we established a collaboration with the VIB BioImaging Core to adapt the workflow for additional light-sheet microscopy datasets. During this collaboration, the modular workflow structure allowed individual processing steps to be selectively combined into the resulting three preprocessing stages, according to experimental requirements. While the original NuMorph implementation also provided access to individual processing stages, adapting the analysis required manual selection and execution of separate commands. In contrast, nf-core/lsmquant represents these processing options as explicit workflow configurations, simplifying the reproducible execution of different preprocessing strategies. We evaluated the generalizability by applying nf-core/lsmquant to different sample types than originally targeted by NuMorph. One example is shown here for the processing of a neuromuscular organoid dataset (Fig. 8).

**Figure 7:**
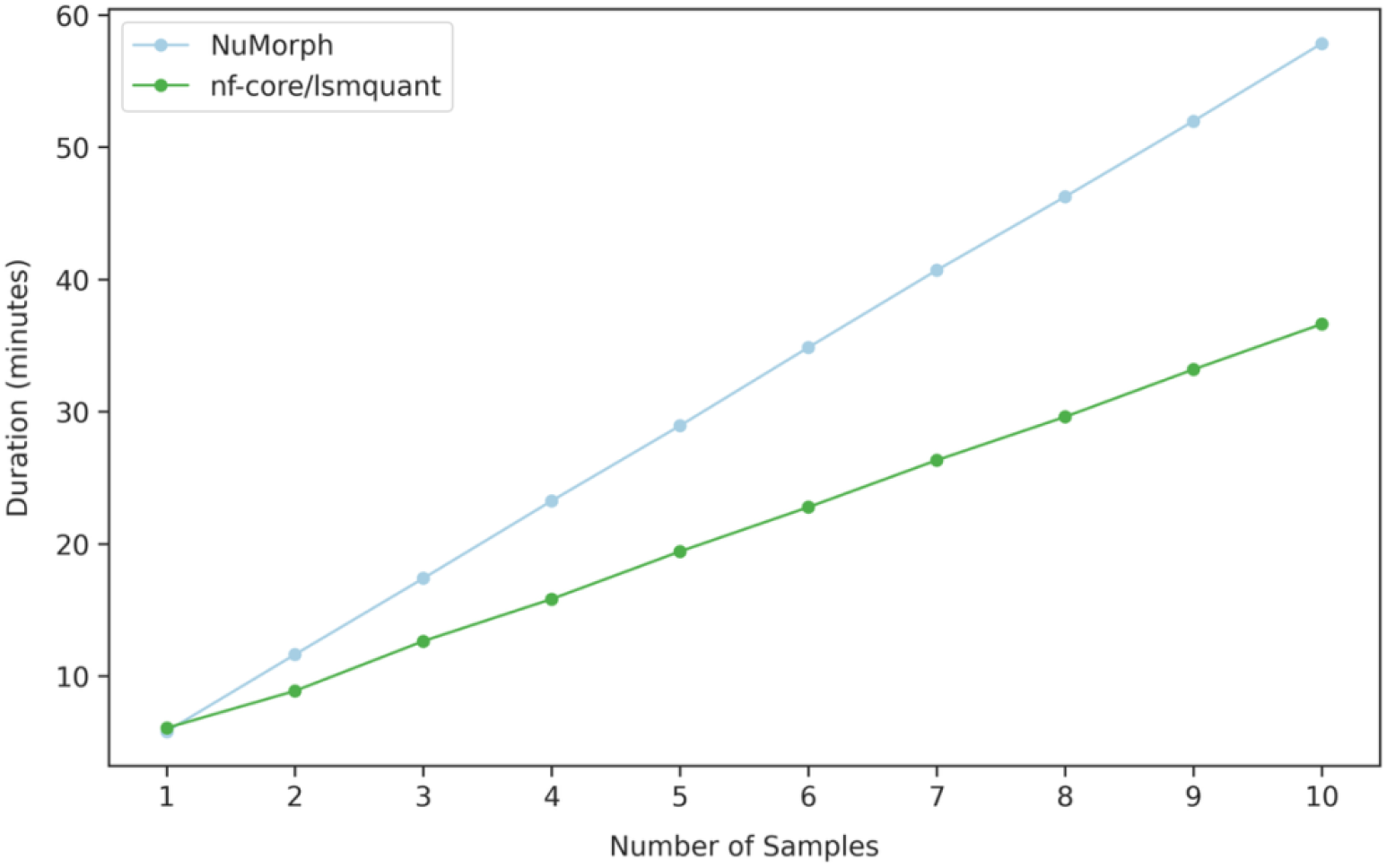
Batch processing comparison of NuMorph and nf-core/lsmquant with increasing sample size. The line plot shows the total wall-clock processing runtime (in minutes) of NuMorph (light blue) and nf-core/lsmquant (green) for increasing number of samples (1–10). Each workflow executed the three preprocessing steps: intensity adjustment, channel alignment, and stitching, and cell nuclei quantification. nf-core/lsmquant consistently requires less time across sample numbers, except for processing 1 sample.

**Figure 8:**
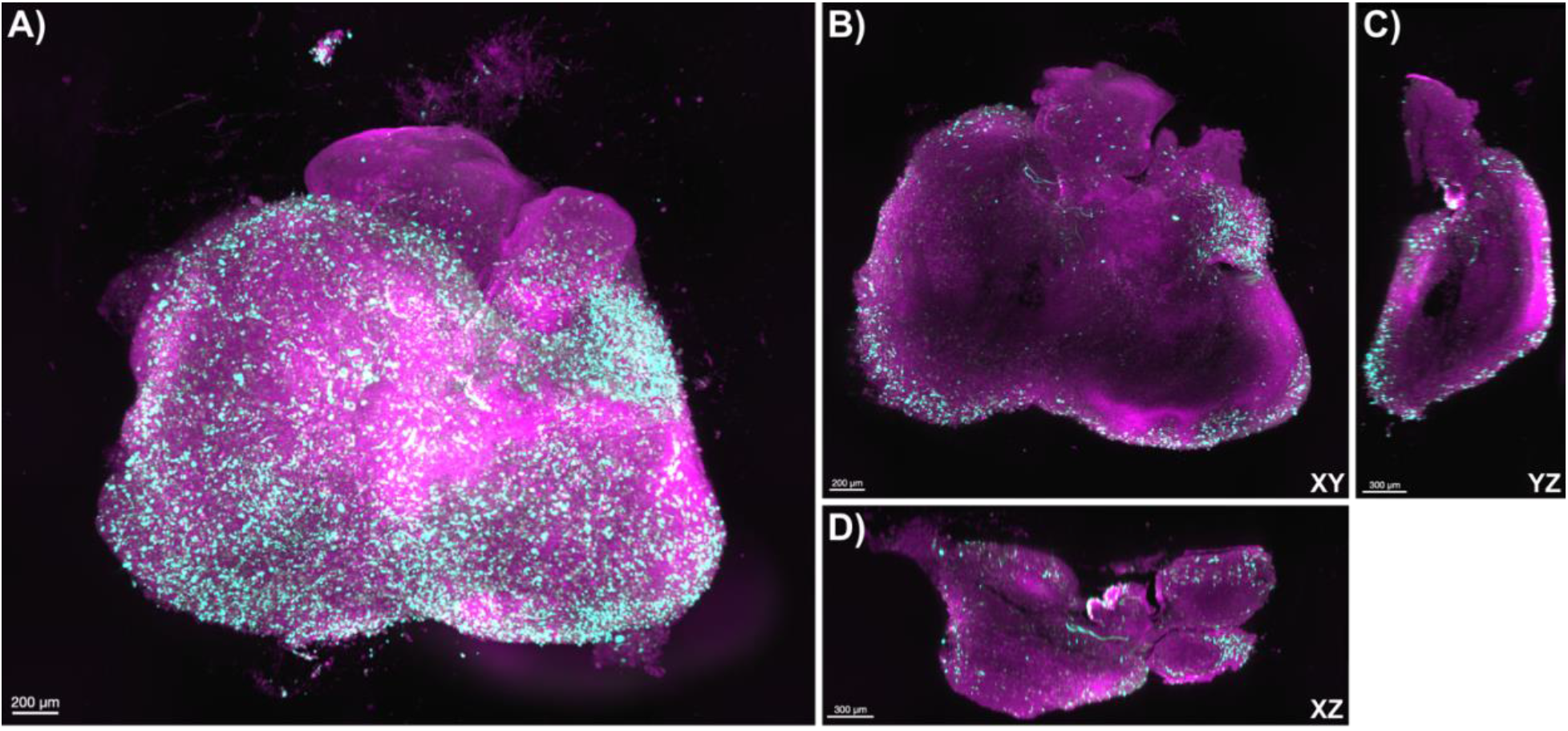
Stitched volume of a neuromuscular organoid processed with nf-core/lsmquant. A shows the 3D volume with two channels: MF20 (purple) and SMI (cyan), of a neuromuscular organoid image, processed with nf-core/lsmquant using the *stitch_only* entry point. Panels B, C, and D show representative 2D planes from different views. The organoid was created in the Ludo Van Den Bosch’s Lab (VIB-KU Leuven Center for Neuroscience) and processed in collaboration with the VIB Bioimaging Core Leuven.

**Figure 9:**
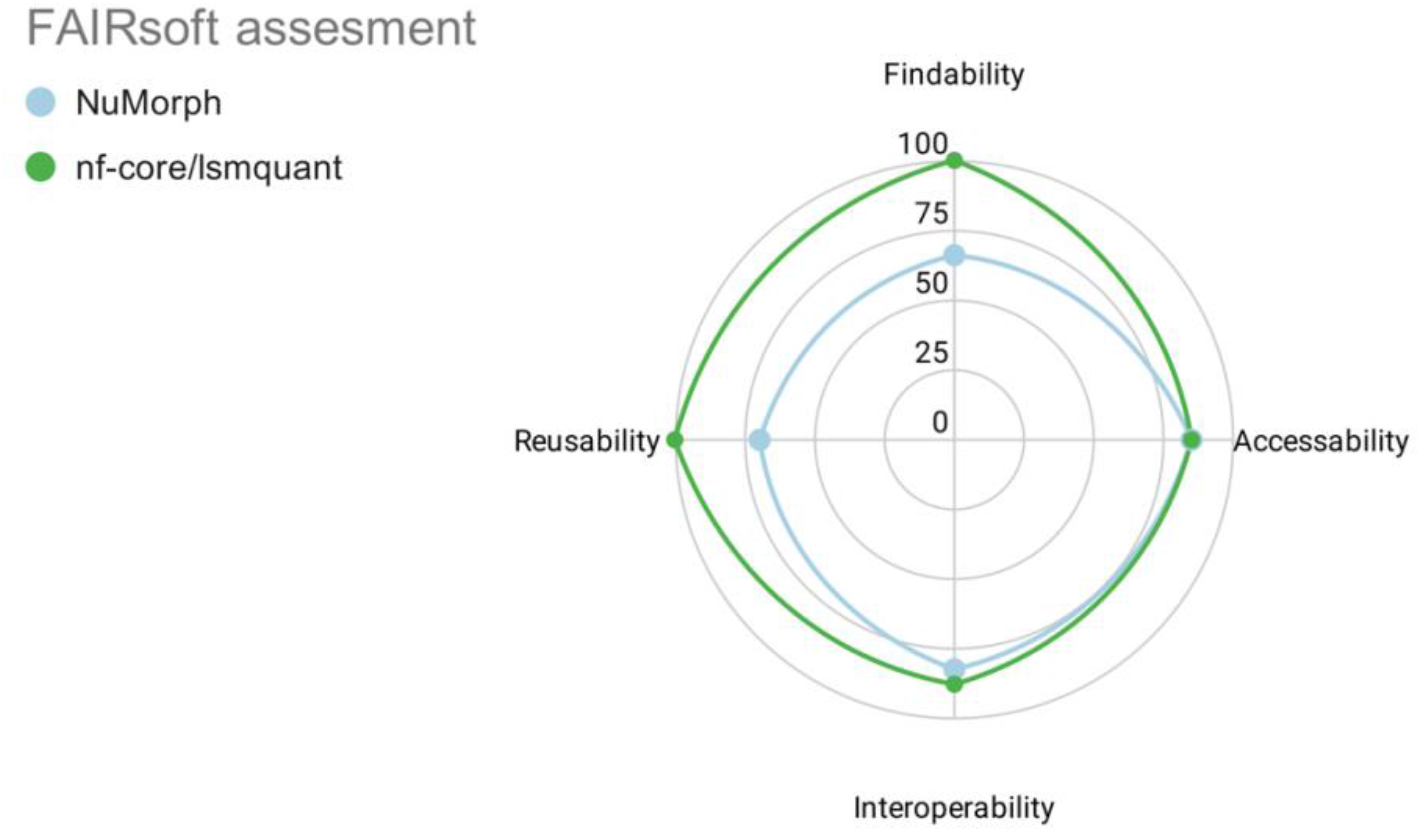
FAIRsoft evaluation of NuMorph and nf-core/lsmquant. Radar plot comparing the FAIRsoft scores of NuMorph (blue) and nf-core/lsmquant (red) across four FAIR indicators: Findability, Accessibility, Properties Interoperability, and Reusability. Scores are expressed as percentages (0–1), with values closer to the outer edge indicating higher compliance with FAIR software principles. nf-core/lsmquant consistently outperforms NuMorph across all four dimensions, reflecting the improved adherence to FAIR principles achieved through the re-engineering process.

The resulting stitched organoid volume showed consistent tile alignment without visible stitching artefacts or discontinuities. Applying nf-core/lsmquant to this additional sample type demonstrates that the workflow can be adapted beyond the original NuMorph development context and provides initial evidence for its applicability to diverse light-sheet microscopy datasets.

## 4. Sustainability evaluation

Finally, we assess how the re-engineering affected software sustainability properties and FAIRness, using the FAIRsoft scoring system[39]. The framework was developed to quantify how research software complies with FAIR principles through hierarchical indicators that represent FAIR software attributes. The detailed scoring for each low and high-level indicator is shown in the supplementary Table 1.

The re-engineering process resulted in a notable increase in overall FAIRness, with NuMorph achieving a total score of 75.9% and nf-core/lsmquant reaching 93.2% (Table S2). Substantial improvements were made in Reusability and Findability, where nf-core/lsmquant achieved scores of 100% in both categories (Table S2). In contrast, NuMorph showed its lowest scores in these areas, with 66% for Findability and 70% for Reusability (Table S2). However, nf-core/lsmquant did not achieve full scores in the categories of Accessibility and Interoperability. The reduced Interoperability score is primarily due to limited flexibility in supported data formats. Both nf-core/lsmquant and NuMorph require input data in a single-channel, single-tile, single z-slice TIFF format for processing and therefore do not fulfill the criterion of flexible data format support.

While the FAIRsoft evaluation provides a structured and quantitative assessment of FAIR-related properties, it does not fully capture additional aspects of software sustainability that are critical for long-term usability and maintainability. Characteristics such as installability, portability, scalability, and extensibility are only partially reflected in FAIR metrics but play a central role in determining whether software can be effectively reused in practice. To complement the quantitative FAIR assessment, we therefore performed a qualitative evaluation of key changes made through our re-engineering process and their impact on software sustainability attributes. These include aspects related to software design, deployment, execution, and maintenance. A structured comparison of the legacy NuMorph implementation and the re-engineered nf-core/lsmquant workflow is summarized in Table 1.

**Table 1:** Comparative summary of NuMorph and nf-core/lsmquant with respect to software sustainability and FAIRness. Overview of key characteristics of the legacy NuMorph toolkit and the re-engineered nf-core/lsmquant pipeline, highlighting the main changes introduced during the re-engineering process and their respective impact on software sustainability and adherence to FAIR software principles.

| Aspect | NuMorph | nf-core/lsmquant | Re-engineering alterations | Impacted Attributes |
| --- | --- | --- | --- | --- |
| Distribution Format | source code | containerized | Compilation and containerization | Portability, Reproducibility |
| Implementation language | MATLAB based | Language agnostic | Nextflow and containerization | Extendibility |
| Software accessibility | Proprietary license for usage | Usage open without proprietary license | Compilation and containerization with the MATLAB runtime | Usability<br>Accessibility<br>Extendibility |
| Installation and dependency management | Source code with partially dependency installation | Automated dependency management one command installation | nf-core best practices, Nextflow, containerization | Installability, Deployability<br>Usability |
| Testing and quality control | No testing | process and workflow tests | nf-core standards and nf-test | Reliability |
| Execution model and parallelization | Manual job submission | Automated process orchestration | Nextflow implementation | Scalability |
| Workflow structure | Nested conditional statements | Workflow management | Nextflow | Understandability, Modularity, Extendibility |
| Documentation | Incomplete and scattered across different sources | Methods description, usage and output information unified | Extended documentation | Understandability |

As shown in Table 1, the re-engineering process improved multiple dimensions of software sustainability attributes. At the level of deployment and execution, the transition from source code distribution to containerized execution environments substantially improves portability and reproducibility. The adoption of Nextflow as the workflow management system enables scalable and parallel execution while removing the dependency on a MATLAB-centric architecture, thereby facilitating the integration of future tools and workflow components irrespective of their implementation language. In terms of usability and accessibility, automated dependency handling and standardized installation and execution procedures reduce the technical barrier for new users.

Specifically, the reimplementation reduces the complexity of running the workflow. The original NuMorph toolkit required users to interact directly with the source code, edit multiple scripts to define parameters and sample information, and execute several commands to start an analysis. For users with limited computational experience, navigating the source code and identifying the relevant configuration points can be challenging. In contrast, the nf-core implementation separates user-provided inputs from the underlying source code through the documented sample sheet and parameter file, while software dependencies are resolved automatically. As a result, pipeline installation and execution can be performed using a single, standardized command without interacting with the source code. Furthermore, by using NuMorph’s processes as compiled standalone applications within Docker, users do not require a proprietary MATLAB License for execution. The introduction of the testing framework nf-test[40] and adherence to nf-core development standards based on continuous integration improves reliability and supports long-term maintenance.

From a software design perspective, the transformation from a monolithic codebase with tightly coupled workflow logic to a modular workflow architecture significantly improves understandability and extensibility. Individual components can now be modified or replaced independently, facilitating the integration of new methods and future development.

## Discussion

Here, we demonstrate that orphaned research software can be systematically rescued and transformed into sustainable, reusable workflows by applying established best practices in scientific software development. By re-engineering the MATLAB-based NuMorph toolkit into the nf-core/lsmquant pipeline, we show how legacy monolithic code can be transformed into a modular, scalable, and reproducible workflow that aligns with community standards and FAIR principles.

The original NuMorph toolkit illustrates a challenge shared by many academic software projects. Although it provided an end-to-end solution for large-scale light-sheet microscopy data processing, the software relied on a single developer and did not achieve broad community adoption. As a result, valuable domain knowledge and implementation effort remained difficult to access and reuse. Factors such as limited discoverability, sparse documentation, and the tight coupling between code and execution environment likely contributed to these barriers. These observations are consistent with broader challenges in research software, where limited adoption of sustainable software development practices, accessibility, and long-term support structures can hinder the reuse of valuable computational methods[41].

The frameworks of FAIR4RS and software sustainability provide complementary perspectives for improving the longevity and reuse of research software. FAIR4RS establishes principles to make software findable, accessible, interoperable, and reusable, while software sustainability focuses on the engineering practices required to maintain and evolve software over time. Our results highlight that these concepts address overlapping but distinct aspects of software reuse. Although NuMorph achieved a comparatively high FAIR assessment score, practical reuse remained challenging due to outdated dependencies, fragmented documentation, and the effort required for installation and execution. This demonstrates that FAIR compliance alone does not necessarily translate into immediate usability for researchers. Instead, effective reuse requires additional technical sustainability attributes, including maintainable software architectures, reproducible execution environments, automated testing, and clear documentation. Therefore, FAIR4RS should be considered an essential foundation for software reuse, while sustainable software engineering practices provide the mechanisms required to preserve and extend that reuse over time.

In this context, an interview study with research software engineers identified several key factors that promote sustainability, including modularity and encapsulation, infrastructure independence, testing, comprehensive documentation, and understandable software[42]. Many of these characteristics are inherently supported by Nextflow and the nf-core framework. Through workflow abstraction with Nextflow, we achieved task-level orchestration. This is beneficial for batch processing multiple samples compared to default sequential NuMorph processing, where we could demonstrate that nf-core/lsmquant requires less overall runtime for processing multiple samples. While NuMorph can theoretically be run concurrently, this requires manual intervention for job submission, restarting failed processes, and awareness of resource allocation. This often tedious work is automatically handled by Nextflow, which simplifies scalability. Furthermore, the integration of other tools into the workflow is language-agnostic, which removes constraints imposed by the tightly coupled, MATLAB-centric architecture and supports the long-term evolution of the workflow.

By embedding the pipeline within the active nf-core community rather than maintaining it as a standalone project, further development and maintenance can be distributed across contributors from multiple research institutions. The nf-core framework reduces maintenance complexity by decomposing workflows into standardized, reusable modules that encapsulate individual bioinformatics tools. This allows contributors to update and maintain individual components independently, while improvements can be efficiently integrated into downstream workflows using nf-core tooling. Consequently, maintenance efforts are distributed across smaller, well-defined tasks rather than concentrated within a single monolithic codebase[43]. In addition, integration into a visible community ecosystem increases the likelihood of attracting new users and contributors, further supporting knowledge transfer and long-term development. Together with continuous integration, automated testing, and standardized code review at both the module and pipeline levels, these practices reduce the risk of software becoming orphaned while ensuring functionality and reliability.

These engineering practices also translate into practical benefits for end users. Separating user-configurable parameters from the underlying source code reduces complexity by not exposing users to it directly and reduces the risk of unwanted modifications. Combined with a standardized execution that remains consistent across local workstations, HPC systems, and cloud environments, lowers the technical barrier to executing complex workflows. Furthermore, aggregating software usage instructions and methodological documentation in a single location improves transparency and makes the workflow easier to understand and adopt.

Our evaluation showed that with the re-engineering process, we were able to preserve NuMorph’s functionalities, and nf-core/lsmquant reproduces the same results at similar computational costs. The slightly longer runtimes we observed for the three processing steps: intensity adjustment, channel alignment, and stitching, can most likely be attributed to the start-up time of the Nextflow process and container. Overall, the differences are negligibly small compared to the overall runtime. Only the new implementation of the cell nuclei quantification process in nf-core/lsmquant differed greatly compared to the legacy NuMorph implementation. With the new implementation, we were able to reduce the runtime by 34.8%, but the peak memory consumption increased by 38.3%. While containerization can preserve and improve access to computational methods, packaging decisions can introduce trade-offs in computational resource requirements.

The practical value of the redesigned workflow was further demonstrated through its adoption by collaborators outside the original development group. In collaboration with the VIB BioImaging Core, the modular architecture could be readily adapted into multiple preprocessing workflows tailored to different experimental requirements. This illustrates how workflow abstraction and development within a community can support adaptation to new applications and collaborative development.

Our work provides a practical strategy for recovering abandoned software and aligning it with principles of FAIR research software and software sustainability. Rather than reimplementing and restructuring the entire legacy code, the reengineering approach follows a pragmatic *strangler fig pattern*[44] that wraps the tightly coupled monolithic codebase into a modular architecture in which NuMorph’s core processes are encapsulated in containerized environments.

Despite these improvements, important challenges remain. The initial re-engineering process itself is technically demanding and requires expertise in both the underlying scientific domain and software engineering practices. Although our approach enables the gradual replacement of legacy components, the current implementation still depends on the original tightly coupled MATLAB-based tools. Consequently, further modernization and maintenance of these components remain challenging due to their complexity. Aside from these architectural constraints, the underlying data model, which is based on collections of individual 2D image files, is not well suited for very large datasets because of the large number of generated files and the associated input/output overhead. Supporting TIFF stacks or next-generation formats such as OME-Zarr, developed by the Open Microscopy Environment (OME), which provide chunked and scalable access to multidimensional image datasets with extensive metadata, represents an important direction for future open source development and improved scalability as dataset sizes continue to grow[45].

In conclusion, we showed that through systematic re-engineering and the adoption of FAIR and software sustainability principles, orphaned research software can be rescued. By transforming the NuMorph toolkit into the nf-core/lsmquant workflow, we preserved valuable domain knowledge while increasing findability, accessibility, and effective reuse. Together, our findings show that combining workflow technologies, community-driven development, and sustainable software engineering practices can preserve valuable research software while improving its accessibility, maintainability, and potential for continued reuse.

## Acknowledgements

We want to thank Robert Prior and Kristel Eggermont from Van Den Bosch’s Lab (VIB-KU Leuven Center for Neuroscience) for sharing the stitched images of their neuromuscular organoid as a demonstration included in the generalization results section. Furthermore, we want to thank the nf-core community, which was helpful throughout the process of creating nf-core/lsmquant.

## Methods

### Reverse engineering

Reverse engineering was performed by using the following resources:

- Original Publication of the NuMorph toolkit[30]
- Wiki page on the Bitbucket repository of NuMorph [46]
- The source code of NuMorph taken from the corresponding GitHub repository[47]
- Source code of NuMorph on BitBucket[48]
- Conversations with the Stein Lab from the University of North Carolina at Chapel Hill where the NuMorph toolkit was developed

Based on these resources, the abstract representation of the NuMorph toolkit and the detailed control flow diagram for the identified core processes were created using the Miro Web tool. Information about the number of files and lines of code was obtained by the command-line tool Cloc version 2.06[49], and language distributions were taken from its GitHub repository About section.

### Modification

Based on the findings collected during reverse engineering, necessary modifications were formulated and applied to the NuMorph toolkit. The source code from NuMorph’s GitHub repository was cloned locally and pushed to the new GitHub repository: qbic-pipelines/Numorph- toolkit[50]. The planned modifications to NuMorph’s source code are summarized in the following table and were applied to the new repository.

| Requirement | Modification |
| --- | --- |
| Distribution of functions without requiring a proprietary MATLAB license | Creation of compiled standalone applications and distribution with the MATLAB Runtime |
| Command-line execution | Implementation of a command-line interface (CLI) |
| User input in open file formats | Define User input in the CLI and MATLAB function to convert it into the data structure required by the toolkit |
| Containerization | For each Matlab stand-alone application, creating a docker image using a Matlab base image with the matlab runtime |
| Output in open file formats | Implementation of a separate tool for the conversion of MATLAB files into CSV or JSON format, `mat2json` |
| Controlled execution of individual processing steps | Implementation of a CLI parameter, controlling execution, by modification in the source code |
| Implementation of a pip-installable package of the 3D U-Net | Separation and restructuring of the 3D U-Net code into a dedicated pip package using a separate repository to facilitate installation and maintenance |
| Functional pip environment for the 3D U-Net | Updating dependency versions to resolve incompatibilities, replacing unavailable packages, and ensuring compatibility with the containerized CUDA toolkit |

After all necessary changes to NuMorph’s source code were implemented, the MATLAB source code was compiled with MATLAB Compiler version 2023a, which corresponds to NuMorph’s original development version to avoid large language incompatibility issues. A GitHub release of the compiled *Numorph-tools* version 1.0.0 was made according to semantic versioning, containing the compiled code for NuMorph’s preprocessing functions, intensity adjustments, channel alignment and stitching, and NuMorph’s analysis functions resample and ara registrations. For each compiled binary, a separate Docker container was created using a MATLAB base image containing the MATLAB runtime version 2023a and installing the corresponding released *Numorph-tools* binaries into the container at build time. The built containers, *numorph_preprocessing* and *numorph_analyze*, were uploaded to the nf-core repository on quay.io. The NuMorph processes intensity adjustment and resampling were added to the nf-core/modules repository together with the corresponding Dockerfiles for creating the final container: numorph/intensity[51] and numorph/resample[52].

The conversion tool *mat2json,* which converts proprietary MATLAB files into the open CSV or JSON file formats, was implemented in MATLAB. The compiled tool version 1.0.0 was released on the GitHub repository qbic-pipelines/mat2json[53] together with the tools’ source code. The tool was added to the nf-core/modules repository as mat2json[54] together with the final Dockerfile, which was used to create and publish the container on the nf-core repository on quay.io. The final container uses a base image containing the MATLAB runtime version 2023a.

The entire Python code, which implements the 3D-Unet-based cell nuclei quantification processes, was separated from the *Numorph-tools* repository and moved to a separate repository in qbic- pieplines/numorph_3dunet[55]. The restructuring resulted in a Python package which is available on the Python Package Index (PyPI): numorph-3dunet[56]. For the final Docker container, we used an NVIDIA-optimized TensorFlow base image: nvcr.io/nvidia/tensorflow:20.10-tf1-py3[57] that has an optimized version of TensorFlow and required CUDA libraries pre-installed. The PyPI package of the NuMorph 3D U-Net was installed via pip during build time, and the resulting container *numorph_3dunet* was uploaded to the nf-core repository on quay.io. The tool is integrated into the nf-core/modules repository as numorph/3duent[58] together with the resulting Dockerfile.

### Forward engineering

The pipeline nf-core/lsmquant was built in Nextflow’s domain-specific language 2 (DSL 2) and the nf-core pipeline template. Using the four built container images, seven Nextflow modules were implemented: *numorphintensity*, *numoprhalign*, *numorphstitch*, *numorphresample*, *numorphregistration*, *numorph3dunet*, and *mat2json*. In addition to the main processes, the nf-core community modules *unzip*, *unzipfiles*, and *multiqc* were added to the pipeline for staging input data and creating a standardized pipeline run report using multiqc. Lastly, we created two additional helper modules, *stagefiles* and *validate_parameters*, which are used to stage raw data in a directory into Nextflow’s work directory and validate the user-adjustable parameter csv file with a provided JSON schema.

Based on the information from reverse engineering the NuMorph toolkit and discussion with our collaborators, three preprocessing workflows were created that can be chosen by the mandatory parameter *stage*:

1. **stitch_only:** performs iterative tile stitching on raw images
2. **int_stitch:** Performs intensity adjustment and iterative tile stitching on raw images
3. **int_align_stitch:** Performs intensity adjustment, channel alignment, and iterative stitching on raw images (set as default).

The ARA-registration and nuclei quantification analysis steps were set as optional analysis steps, which can be specified using the respective parameters *–nuclei-quantification and –ara- registration*. All MATLAB output files (.mat) created by each process are collected and converted by the module *mat2json*. The testing framework nf-test was used to implement tests for all newly created modules and workflow-level tests for continuous integration. For the creation of the nf-core/lsmquant pipeline metromap, Inkscape and the nf-core SVG style sheet were used, which provides predefined pipeline components.

### Benchmark experiment using the small test dataset

To evaluate functional equivalence and resource usage of the new system nf-core/lsmquant compared to the legacy system NuMorph, we performed a benchmark experiment on the small test dataset from the original NuMorph Repository on Bitbucket. The small test dataset consists of a subset of a tissue-cleared mouse brain, imaged with the Ultramicroscope II microscope from Lavision (now Miltenyi). The subset was taken from z-plane 0600 to 0620 of 4 tiles from two imaged channels: ctip2 (for deeper-layer neurons) and topro (TO-PRO3 for all nuclei), with a total number of 169 images and a size of 1.39 GiB. This dataset was deposited on Zenodo[59]. On this dataset, we ran nf-core/lsmquant version 1.0.3[33] and the native NuMorph implementation[60] 30 times, with the same parameter settings, including the following four processing steps: intensity adjustment, channel alignment, stitching, and cell nuclei quantification (reported as count). For the experiment, we created a bash script: benchmark.sh[60] to automate and randomize the execution order between nf-core/lsmquant and NuMorph runs. The experiment was performed on a Linux-based workstation with 16 CPU cores, 64 GiB RAM, and 1 NVIDIA GeForce RTX 3090 GPU with 24 GiB RAM.

### Resource usage and runtime comparison

We compared resource usage and runtime of the processes: intensity adjustment, channel alignment, stitching, and cell nuclei quantification of the nf-core/lsmquant implementation and the legacy NuMorph implementation. Runtime and resource usage were captured during the benchmark run on the small test dataset. For nf-core/lsmquant, these values are automatically captured by Nextflow and reported for each process in the .command.trace file in the workflow’s work directory. The pipeline was executed using a config file (benchmark.config[60]) that sets resource boundaries to the host system’s maximum resources available. This is done to ensure that the processes are not restricted through other module or pipeline configurations.

To obtain comparable runtime and resource usage results for the native NuMorph implementation, we adapted Nextflow’s monitoring logic and created a bash script (monitoring_script.sh[60]) that starts NuMorph processing steps. The captured values are reported in a matlab_trace.log file for each process. To compare each processing step’s runtime and resource usage, NuMorph’s processing steps were executed sequentially in a predefined order: intensity adjustment, channel alignment, stitching, and nuclei quantification. One complete sequence of these stages was defined as one complete workflow run.

Evaluation of captured runtimes and resource usage values was performed using the Jupyter notebook resource_and_scaling_results.ipynb[61]. The monitored resources were extracted from the corresponding trace files for each process per replicate workflow run, aggregated into a combined data frame, and raw values were stored in a benchmark.csv. Raw values were converted into human-readable units (milliseconds to seconds and bytes to gigabytes), and mean and standard deviations were calculated per process for each workflow implementation. Wall clock runtime and peak RSS were plotted as barplots, where the height of the bar shows the corresponding mean value and error bars display the 95% interval of datapoints, ranging from 2.5 to the 97.5 percentiles.

### Functional equivalence comparison

To demonstrate the functional equivalence of nf-core/lsmquant to the original NuMorph toolkit, the outputs produced during the benchmark experiment with the small dataset were used and compared using the Jupyter notebook output_comparison.ipynb[61]. For each of the 30 runs per workflow, all pairwise mean squared error (MSE) values of the stitched image volumes for both channels were calculated and visualized in a 30 x 30 heatmap. For the qualitative example, randomly selected replicates from the benchmark experiment for NuMorph (replicate 8) and nf- core/lsmquant (replicate 21), the middle z-plane (z0009) was visualized along with the difference map showing the subtracted intensities and the computed MSE for both channels. The example z-slices are displayed in greyscale using a contrast stretch to the 1st -99th percentile of the NuMorph stitched example image to visualize brightness differences compared to the nf-core/lsmquant stitched image and to improve structure visibility.

We also tested functional equivalence on a tissue-cleared, postnatal day 4 (P4) mouse brain (Sample L73D766P9), which was generated and imaged as previously described by our collaborators from the Lab of Jason L. Stein at UNC at Chapel Hill[37,62]. Briefly, the P4 mouse brain was tissue-cleared with iDISCO+ protocol and immunolabeled with rabbit anti-Brn2 (Cell Signaling Technology, 12137, 1:100) and rat anti-Ctip2 (Abcam, ab18465, 1:400) as the primary antibodies for upper layer cortical neurons and deep layer cortical neurons respectively and goat anti-rat Alexa Fluor 568 (Thermo Fisher, A11077, 1:200) and goat anti-rabbit Alexa Fluor 790 (Thermo Fisher, A11369, 1:50) as the secondary antibodies. The sample was incubated with TO-PRO-3 (Thermo Fisher, T3605, 1:400) dye to label all nuclei in the mouse brain. The sample was then imaged intact using the Ultramicroscope II microscope. The data of the imaged mouse brain L73D766P9 is deposited at the Brain Observatory Storage Service & Database[38,63].

We ran the preprocessing stage (intensity adjustment, channel alignment, and stitching) of nf- core/lsmquant and NuMorph on the P4 mouse brain dataset with identical parameter settings. The middle z-plane 0507 was chosen as a qualitative example of stitched images for the channels: topro, ctip2, brn2. Grayscale images are contrast stretched to the 1st–99th percentile of the NuMorph output, and the same intensity limits are applied to the corresponding nf-core/lsmquant image, to visualize relative brightness differences. For each channel, stitched 2D images of NuMorph and nf-core/lsmquant are subtracted and visualized as pixel-wise difference maps, and the corresponding MSE was calculated. Furthermore, we calculated the MSE between each individual stitched z-plane of NuMorph and nf-core/lsmquant per channel and visualized it as scatter plots to show differences for the complete dataset.

### Scalability of batch processing

To assess the scalability of batch processing, the small test dataset was used to simulate processing multiple samples, starting with one up to ten independent samples as batches with both NuMorph and nf-core/lsmquant. For the legacy implementation, a custom automation script, scaling.sh[60], was developed to execute the individual NuMorph processing steps in the predefined sequence: intensity adjustment, channel alignment, stitching, and nuclei quantification for multiple samples. For nf-core/lsmquant, the same datasets were processed by defining the corresponding samples in the samplesheet. Both workflows were run with the same parameter settings. Total batch processing time (wall-clock time) was recorded for each batch size and used to compare the scalability of the two implementations. For nf-core/lsmquant, the total runtime for a batch was taken from the execution report HTML file after batch completion. For NuMorph, the runtimes of all processes were taken from the matlab_trace.log files and added for the total runtime of each batch.

### Real-world deployment and adoption

To evaluate the real-world applicability of nf-core/lsmquant, the workflow was deployed in collaboration with the VIB BioImaging Core (Leuven, Belgium). A 20 GB light-sheet microscopy dataset of a neuromuscular organoid provided by the Van Den Bosch Lab (VIB-KU Leuven Center for Brain & Disease Research) was processed using the `--stitch_only` workflow of nf-core/lsmquant. The sample was acquired on a Zeiss Lightsheet 7 microscope with a voxel size of 0.93 µm × 0.93 µm × 4.05 µm (x, y, z) and consisted of four image tiles arranged in a 2 × 2 grid, 354 z-slices, and three imaging channels. For stitching, two channels (channel 1: MF20; channel 2: SMI) were selected. The stitched image was converted to the Imaris file format using the Imaris File Converter (Oxford Instruments) and subsequently inspected in Imaris Viewer. Representative orthogonal views were generated from the xy (plane 658), yz (plane 1776), and xz (plane 1319) orientations to assess the quality of the stitched volume. For display purposes only, the minimum intensity threshold of the channel MF20 was set to 184,18 to suppress background signal and improve visual contrast.

### Sustainability evaluation and FAIR assessment

We assessed the FAIRness of both pipelines manually, using the FAIRsoft[39] scoring schema in Excel. The detailed evaluation of individual high- and low-level properties and scoring is shown in the supplementary Table 1 together with the original Excel file. Only the high- and low-level properties for all and non-web-based software were applicable and used for scoring. Scores for each FAIR indicator and the overall score were computed and converted into percentages. For NuMorph and nf-core/lsmquant, the scores of each FAIR indicator were displayed as a radar plot using Excel.

### Availability of supporting source code and requirements

**Source code:**

Project name: nf-core/lsmquant[33]

Project home page: https://github.com/nf-core/lsmquant

Operating system(s): Platform independent

Programming language: Nextflow

Other requirements: Nextflow >= 25.10.4, Docker or Singularity or Apptainer

License: MIT.

Project name: Numorph-tools [50]

Project home page: https://github.com/qbic-pipelines/Numorph-toolkit

Project Container(s):

- https://quay.io/repository/nf-core/numorph_preprocessing [64]
- https://quay.io/repository/nf-core/numorph_analyze [65]

Operating system(s): Linux, Windows, macOS

Programming language: MATLAB

Other requirements: MATLAB version R2023a

License: MIT

Project name: NuMorph-3DUnet [56]

Project home page: https://github.com/qbic-pipelines/numorph_3dunet

Project package: https://pypi.org/project/numorph-3dunet/

Project Container(s): https://quay.io/repository/nf-core/numorph-3dunet [55,66]

Operating system(s): Linux

Programming language: Python

Other requirements: Python 3.6.10, tensorflow-gpu 1.12.0

License: MIT

Project name: mat2json [53]

Project home page: https://github.com/qbic-pipelines/mat2json

Project Container(s): https://quay.io/repository/nf-core/mat2json [67]

Operating system(s): Linux, Windows, MacOS

Programming language: MATLAB

Other requirements: MATLAB

R2023a License: MIT

Project name: rescue_orphan_code_results [61]

Project home page: https://github.com/CaroAMN/rescue_orphan_code_results

Operating system(s): Linux, Windows, macOS

Programming language: Python, Bash

Other requirements:

License: MIT

Project name: NuMorph-orphan-bioinformatics-workflows [60]

Project home page: https://github.com/CaroAMN/NuMorph-orphan-bioinformatics-workflows

Operating system(s): Linux, macOS

Programming language: MATLAB

Other requirements: MATLAB R2023a

License: MIT

### Supporting data

NuMorph test dataset: https://doi.org/10.5281/zenodo.14916478 [59]

NuMorph trained model: https://doi.org/10.5281/zenodo.16893708 [68]

Benchmark results: (in progress)

Original data of the P4 mouse brain sample L73D766P9: https://bossdb.org/project/curtin2026 [38]

Stitched data P4 mouse brain sample L73D766P9: Bioimage archive (in progress) Stitched data neuromuscular organoid: Bioimage archive (in progress)

### List of abbreviations

RSS: Resident Set Size
GiB: Gigabyte
FAIR: Findable, Accessible, Interoperable, Reusable
CPU: Central Processing Unit
GPU: Graphics Processing Unit
RAM: Random Access Memory
CWL: Common Workflow Language
TIFF: Tagged Image File Format
FAIR4RS: Findable, Accessible, Interoperable, Reusable for research software
CSV: Comma-separated values
JSON: JavaScript Object Notation
ARA: Allen Reference Atlas
DSL 2: Domain Specific Language 2
CLI: Command Line Interface
OME-Zarr: Open Microscopy Environment Zarr
MiB: Megabyte
GUI: Graphical User Interface
ZIP: Compressed File Type (.zip)
PyPI: Python Package Index
P4: postnatal day 4

### Declarations

#### Ethics statement

In this study, no new animal experiments were performed, and no new animal data were collected. The mouse brain datasets used for evaluation were previously generated by our collaborators in the laboratory of Jason L. Stein at the University of North Carolina at Chapel Hill and were obtained from the associated published studies[37,38]. Ethical approval for the original animal experiments was obtained by the investigators who generated the datasets, and all animal procedures were conducted in accordance with the relevant institutional and regulatory requirements.

#### Authors’ contributions

Individual authors’ contributions are listed below based on the CRediT Contributor Roles Taxonomy.

Carolin Schwitalla: Software, Writing original draft, Conceptualization, Investigation, Visualization, Formal Analysis, Project administration, Validation

Luis Cullar Kuhn: Funding Acquisition, Writing – review & editing, Supervision, Project administration Matthias Hörtenhuber, Software

Niklas Grote, Writing – review & editing, Software, Investigation Tatiana Woller, Writing – review & editing, Software, Investigation

Irene Lamberti, Writing – review & editing, Software, Investigation, Resources Benjamin Pavie, Writing – review & editing, Software, Investigation

Thomas Küstner, Writing – review & editing, Supervision

Felix A. Kyere, Writing – review & editing, Investigation, Resources, Project administration Ian Curtin, Writing – review & editing, Resources, Project administration

Jason L. Stein: Funding Acquisition, Writing – review & editing, Project administration

Sven Nahnsen: Funding Acquisition, Writing – review & editing, Conceptualization, Supervision, Project administration

#### Disclosure of use of AI-assisted tools including generative AI

In preparing the manuscript, generative AI tools were used to assist with text refinement and language editing. The specific tools are: Grammarly, ChatGPT Free plan (models: GPT-4o mini, GPT-4o, GPT-5 variants), Claude Free plan (models: Sonnet 4.6).

