## Supplemental Figure 1 for "From Abandoned Scripts to FAIR Community Pipelines: Rescuing Orphan Bioinformatics Workflows with nf-core — Lessons from Light-Sheet Fluorescence Microscopy"

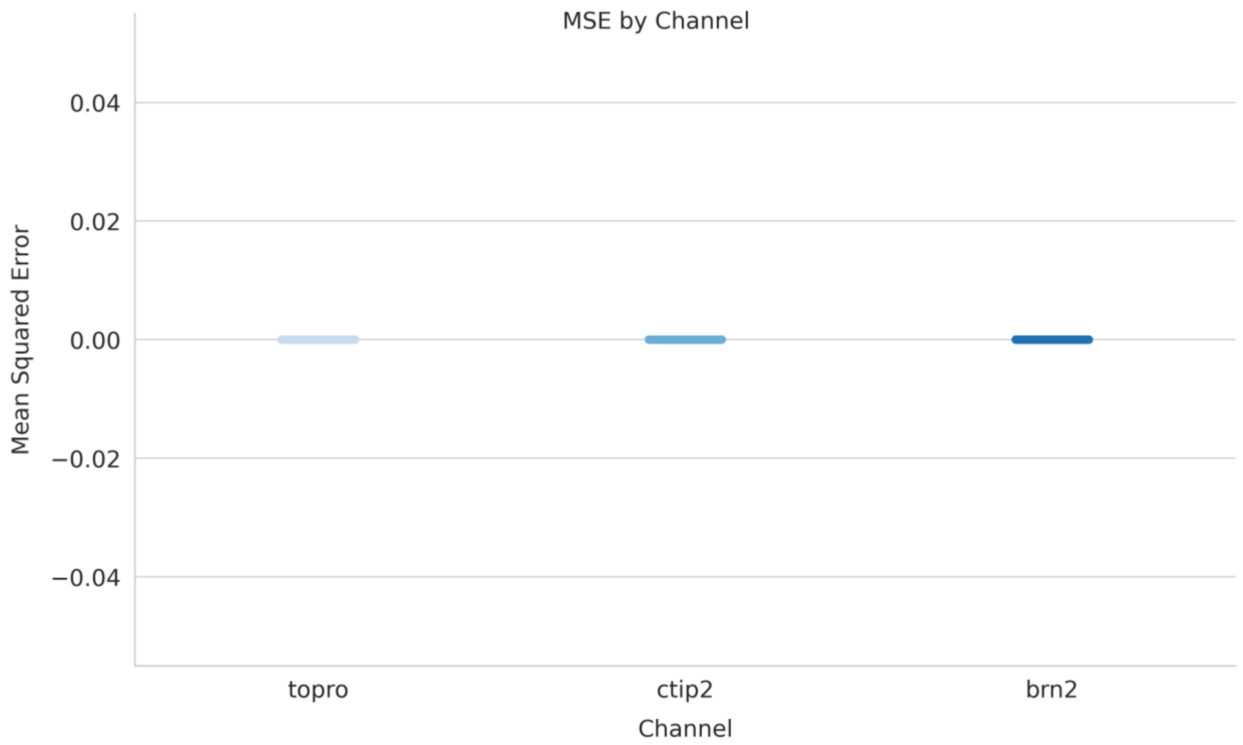


**Supplementary Figure 1: Per z-plane image differences between NuMorph and nf-core/lsmquant outputs on the large mouse brain dataset.** Mean squared error (MSE) was computed between corresponding stitched z-planes produced by NuMorph and nf-core/lsmquant for each fluorescence channel (TO-PRO, Ctip2, Brn2; n=1,015 z-planes per channel). Each point represents one z-plane comparison. MSE values are zero across all planes and all channels, demonstrating pixel-level equivalence between the two pipeline outputs on the large mouse brain dataset.
