## Supplemental Table 1 for "From Abandoned Scripts to FAIR Community Pipelines: Rescuing Orphan Bioinformatics Workflows with nf-core — Lessons from Light-Sheet Fluorescence Microscopy"

**Supplementary Table 1: Summary statistics for the benchmarking experiments comparing the original NuMorph toolkit and the nf-core/lsmquant workflow**. For each processing step, the mean and standard deviation of runtime and peak resident set size (RSS) across all benchmark runs are reported.

| | | Runtime [minutes] | | Peak RSS [GiB] | |
|---|---|---|---|---|---|
| | | mean | std | mean | Std |
| NuMorph | Intensity | 0.5594505556 | 0.0082518249 | 1.5484003703 | 0.031592647 |
| | Alignment | 1.44105 | 0.0166182244 | 1.4613422394 | 0.0718654553 |
| | Stitching | 1.2857633333 | 0.0137498864 | 1.7782979329 | 0.062061124 |
| | Count | 2.5068955556 | 0.036266925 | 10.3759553274 | 0.0179188374 |
| nf-core/lsmquant | Intensity | 0.5801827778 | 0.0052817358 | 1.4283356984 | 0.0283437524 |
| | Alignment | 1.4987822222 | 0.0143120759 | 1.4789169312 | 0.0215403878 |
| | Stitching | 1.3563844444 | 0.0155786227 | 1.7505407969 | 0.0348294065 |
| | Count | 1.6337811111 | 0.0087997646 | 14.3594420115 | 0.0165110496 |
| | | Absolute mean difference [seconds] | | Absolute mean difference [MiB] | |
| | Intensity | 1.2439332 | | 122.94622208 | |
| | Alignment | 3.4639332 | | 17.99648256 | |
| | Stitching | 4.2372666 | | 28.42331136 | |
| | Count | 52.3868664 | | 3,98348668 [GiB] | |
