## Supplemental Table 2 for "From Abandoned Scripts to FAIR Community Pipelines: Rescuing Orphan Bioinformatics Workflows with nf-core — Lessons from Light-Sheet Fluorescence Microscopy"

**Supplementary Table 2: Indicator-level FAIRsoft assessment of the original NuMorph software and the nf-core/lsmquant reimplementation**. Scores are reported for all FAIRsoft indicators and aggregated FAIR dimensions (Findability, Accessibility, Interoperability, and Reusability).

| **Findability** | | Low level indicators | | Weight | NuMorph | NuMorph score | nf-core/lsmquant | nf-core/lsmquant score |
|---|---|---|---|---|---|---|---|---|
| Identity uniqueness | F1.1 | Uniqueness of name | 0,8 | 0,4 | TRUE | 0,32 | TRUE | 0,4 |
| | F1.2 | Identifiability of version | 0,2 | | FALSE | | TRUE | |
| Existence of metadata | F2.1 | Existence of structured metadata | 0,6 | 0,2 | FALSE | 0 | TRUE | 0,2 |
| | F2.2 | Existence of standardized metadata | 0,4 | | FALSE | | TRUE | |
| Discoverability | F3.1 | Discoverabilities in software registries | | 0,4 | FALSE | 0,34 | TRUE | 0,4 |
| | F3.2 | Discoveries in software repositories | | | TRUE | | TRUE | |
| | F3.3 | Discoverability in literature | | | TRUE | | TRUE | |
| | | | | | | 0,66 | | 1 |
| **Accessibility** | | | | | | | | |
| Existence of an available working version | A1.1 | Existence of an API or web interface | 0,6 | 0,7 | X | 0,7 | X | 0,7 |
| | A1.2 | Existence of a downloadable and buildable software working version | 0,5 | | TRUE | | TRUE | |
| | A1.3 | Existence of installation instructions | 0,2 | | TRUE | | TRUE | |
| | A1.4 | Existence of test data | 0,1 | | TRUE | | TRUE | |
| | A1.5 | Existence of software source code | 0,2 | | TRUE | | TRUE | |
| Software history trackability | A2.1 | Existence of metadata of previous versions in software repositories | 0 | 0 | FALSE | 0 | TRUE | 0 |
| | A2.2 | Existence of accessible previous versions of the software | 0 | | FALSE | | TRUE | |
| Unrestricted access | A3.1 | No registration is required | 0 | 0,3 | FALSE | 0,15 | TRUE | 0,15 |
| | A3.2 | Availability of a version for an open-source operating system | 0,25 | | TRUE | | TRUE | |
| | A3.3 | Availability for several OS | 0,25 | | TRUE | | TRUE | |
| | A3.4 | Availability through publicly available e-infrastructures | 0,25 | | FALSE | | FALSE | |
| | A3.5 | Availability through several e-infrastructures | 0,25 | | FALSE | | FALSE | |
| | | | | | | 0,85 | | 0,85 |
| **Interoperability** | | | | | | | | |
| Data format standards and practices | I1.1 | Use of standard data formats | 0,5 | 0,6 | TRUE | 0,48 | TRUE | 0,48 |
| | I1.2 | Use of standard API specification framework | 0 | | X | | X | |
| | I1.3 | Verifiability of data formats | 0,3 | | TRUE | | TRUE | |

| | | | | | | | | |
|---|---|---|---|---|---|---|---|---|
| | I1.4 | Flexibility of data format support | 0,2 | | FLASE | | FALSE | |
| | I1.5 | Generation of provenance information | 0 | | FALSE | | TRUE | |
| Software integration | I2.1 | Existence of API/Library /version | 0,5 | 0,1 | FALSE | 0,05 | TRUE | 0,1 |
| | I2.2 | E-infrastructure compatibility | 0,5 | | TRUE | | TRUE | |
| Dependencies availability | I3.1 | Existence of dependence statements | 0,33 | 0,3 | TRUE | 0,297 | TRUE | 0,297 |
| | I3.2 | Availability of software dependencies | 0,33 | | TRUE | | TRUE | |
| | I3.3 | Availability through dependencies-aware systems | 0,33 | | TRUE | | TRUE | |
| | | | | | | 0,827 | | 0,877 |
| **Reusability** | | | | | | | | |
| Existence of Usage Documentation | R1.1 | Existence of usage guides | 1 | 0,3 | TRUE | 0,3 | TRUE | 0,3 |
| | R1.2 | Existence of usage examples | 0 | | TRUE | | TRUE | |
| Existence of license and or terms of use | R2.1 | Existence of terms of use | 1 | 0,3 | X | 0 | X | 0,3 |
| | R2.2 | Existence of conditions of use | 1 | | FALSE | | TRUE | |
| Existence of contribution recognition and governance | R3.1 | Existance of contributionrs policies | 0 | 0,2 | FALSE | 0,2 | TRUE | 0,2 |
| | R3.2 | Existence of credit | 1 | | TRUE | | TRUE | |
| Existence of versioning and history traceability | R4.1 | Use of version control | 1 | 0,2 | TRUE | 0,2 | TRUE | 0,2 |
| | R4.2 | Existence of a release policy | 0 | | FALSE | | TRUE | |
| | R4.3 | Existence of metadata of previous versions in software repositories | 0 | | FALSE | | TRUE | |
| | | | | | | 0,7 | | 1 |
| | | Total | | 4 | | 3,037 | | 3,727 |
| | | Percentage | | 100 | | 75,925 | | 93,175 |
| All software | | | | | | | | |
| non web based | | | | | | | | |
| web based | | | | | | | | |
